# The distinct phenotypic signatures of dispersal and stress in an arthropod model: from physiology to life history

**DOI:** 10.1101/581116

**Authors:** Maxime Dahirel, Stefano Masier, David Renault, Dries Bonte

## Abstract

Dispersing individuals are expected to encounter costs during transfer and in the novel environment, and may also have experienced stress in their natal patch. Given this, a non-random subset of the population should engage in dispersal and eventually show divergent stress-related responses towards new conditions. Dispersal allows escape from stress, but is equally subjecting individuals to it.

Physiological shifts expressed in the metabolome form a major part of responses to stress exposure and are expected to be associated with the dispersal phenotype, thereby shaping physiological dispersal syndromes. We analyzed how metabolic profiles and life-history traits varied between dispersers and residents of the model two-spotted spider mite *Tetranychus urticae*, and whether and how these syndromes varied with exposure to a stressful new host plant (tomato). Regardless of the effect of host plant, we found a physiological dispersal syndrome where, relative to philopatric individuals, dispersers were characterized by lower leaf consumption rates and a lower concentration of several amino acids, indicating a potential dispersal-foraging trade-off. As a possible consequence of this lower food intake, dispersers also showed a lower reproductive performance. Responses to tomato exposure were consistent with this plant being a stressor for *Tetranychus urticae*, including reduced fecundity and reduced feeding by mites. Tomato-exposed mites laid larger eggs, which can be interpreted as a plastic response to food stress, increasing the likelihood of survival to maturity. Contrary to what could be expected from the costs of dispersal and stress resistance and from previous meta-population level studies, there was no interaction between dispersal status and host plant for any of the examined traits, indicating that the impacts of a new stressful host plant are equally incurred by residents and dispersers.

We thus provide novel insights in the processes that shape dispersal and the putative feedbacks on ecological dynamics in spatially structured populations.

## Introduction

Dispersal, i.e. movement leading to gene flow, is a key trait at the nexus between ecological and evolutionary dynamics (Bonte and Dahirel, 2017; Clobert et al., 2012; Govaert et al., 2019). Not all individuals or populations show the same dispersal motivation and ability. The realization that this variation is often non-random has led to the study of dispersal syndromes, i.e., environmentally- and/ or genetically-driven correlations between dispersal propensity/ability and other traits (life history, behavior, morphology…), and their consequences (Bonte and Dahirel, 2017; Cote et al., 2017; Ronce and Clobert, 2012). Among other things, dispersal syndromes may arise because dispersing and non-dispersing individuals actually experience different environmental and social contexts throughout their life, and thus different selective pressures favoring different trait combinations (Ronce and Clobert, 2012).

As their environment varies in space and time, organisms encounter a variety of conditions during their lifetime, and may be forced to deal with unfavorable or stressful conditions (Steinberg, 2012). A large corpus of research has shown that a mosaic of behavioral, physiological or biochemical responses are triggered during exposure to mildly to harshly stressful environmental conditions (Steinberg, 2012; Sulmon et al., 2015; Tuomainen and Candolin, 2011), with potential side-effects on other individual life traits and fitness (Telonis-Scott et al., 2006). Individuals from the same population can show a large variation in stress resistance, which can be of both environmental (e.g. Henry et al., 2018; Ximénez-Embún et al., 2016) and genetic origin (Gerken et al., 2015; Rion and Kawecki, 2007; Rolandi et al., 2018).

Taking that into account, the relationship between dispersal and the ability of organisms to cope with environmental stressors is potentially complex. One could expect stress-sensitive individuals to increase their emigration rates under stress, to limit the negative fitness consequences of stress; this would lead to a negative correlation between dispersal tendency and stress resistance. A negative correlation between dispersal and stress resistance may also arise if stress resistance is costly, as costs incurred during dispersal (Bonte et al., 2012) may limit dispersers’ future stress resistance ability or vice versa (Matsumura and Miyatake, 2018). In all cases however, the balance between fitness expectations at ‘home’ versus elsewhere will determine the eventual dispersal strategy. This means relationships between stress resistance and dispersal can be equally driven by fitness expectations after immigration in novel environments (Renault et al., 2018). Traits or contexts favoring dispersal may also favor stress tolerance (Lustenhouwer et al., 2019). As the probability of arriving in marginal/suboptimal habitat should increase in heterogeneous environments, stress-resistant individuals may eventually profit most from dispersal, giving rise to positive correlations between these traits.

The variation of physiological responses in relation to the environment and phenotype can be explored using metabolomic approaches (Bundy et al., 2008). As for morphological, behavioral and life-history traits (Beckman et al., 2018; Cote et al., 2017; Ronce and Clobert, 2012), information about whole-organism physiological state variation should help us understand the underlying mechanisms shaping dispersal variability. Metabolic profile changes in response to stress exposure, and their (putative) functional roles, have been extensively documented, especially in arthropods (Bundy et al., 2008; Steinberg, 2012; Teets and Denlinger, 2013). In contrast, metabolite differences between dispersers and residents even in interaction with the environment- are surprisingly understudied, compared to other phenotypic differences (Cote et al., 2017; Ronce and Clobert, 2012; but see Tung et al., 2018; Van Petegem et al., 2016).

Using the phytophagous two-spotted spider mite *Tetranychus urticae* Koch 1836, we combined metabolic profiling to more classical life-history and morphological measurements, and investigated whether and how dispersing and philopatric individuals differed from each other and in their resistance to a biotic stressor. We expected dispersing and philopatric individuals to differ in several traits (i.e. to exhibit dispersal syndromes), with dispersers in particular showing detectable evidence of incurred dispersal costs, such as lower fecundity (Matsumura and Miyatake, 2018). *Tetranychus urticae* is highly generalist at the species level, with over 1100 documented host plant species (Migeon et al., 2010). There is, however, strong among-strain variation in the ability to survive and reproduce on specific host plants (reviewed in Rioja et al., 2017). In this species, metapopulation structures that favor the evolution of lower dispersal capacity or delayed dispersal also lead to the evolution of higher performance on a new stressful host (De Roissart et al., 2015; De Roissart et al., 2016). This implies that dispersal and (biotic) stress resistance are negatively correlated, for instance because dispersers are stress-intolerant individuals that flee adverse conditions when they can, finding themselves at a fitness disadvantage if they cannot. However, this link was not formally established, and the relationship between dispersal and stress tolerance may have arisen instead due to third factors, such as density-dependent competition (Bonte et al., 2014; De Roissart et al., 2015; De Roissart et al., 2016). We here more formally test this hypothesis at the individual level: although an overall performance decrease was expected in all stressed mites, we predicted that the metabolic and life-history profiles of dispersers would be more negatively influenced by exposure to the new stressful host plant than those of residents.

## Material and methods

### Mite strain and rearing

We used mites from the bean-adapted LS-VL strain, which has been maintained on whole bean plants since 2000 (*Phaseolus vulgaris* L. cv. Prélude) (De Roissart et al., 2016; Van Leeuwen et al., 2004). Prior to experiments, synchronized mites of known age were obtained by letting mites from the stock population lay eggs for 24h on 7 × 7 cm^2^ bean leaf squares (freshly cut from two-week-old plants) set on wet cotton (50 mites/ leaf square), then removing them. These eggs were then maintained at 30 °C, L:D 16:8 until reaching adulthood (i.e. ~8-10 days), with the cotton kept hydrated. All adult mites used in subsequent experiments resulted from this procedure.

### Dispersal trials

We used two-patch setups to sort dispersers from resident mites. One- to two-day-old mated adult females (the main dispersive stage; Krainacker and Carey, 1990) were placed on freshly cut 4 cm^2^ bean leaf squares (start patches), which were then each connected to a 4 cm^2^ empty bean leaf square (target patch) using a Parafilm bridge (2 × 8 cm^2^). Seventy bridges were used across all experiments. One hundred mites were initially placed on each start patch (25 mites.cm^−2^, an intermediate density in the range tested by Bitume et al., 2013). To keep leaves hydrated and limit mite escapes, bridges were set on wet cotton and external edges covered with 2 mm wide strips of moist tissue paper. Mites were then let free to disperse for 48h at 25 °C, L:D 16:8; individuals found on the target patch were then deemed dispersers, individuals found on the start patch considered residents. Detailed records of disperser numbers were only kept for the first 10 out of 70 bridges. On these, 18.5 % of mites successfully dispersed (95% confidence interval: 16.2 - 21.0 %; range: 5 - 32 %).

### Experiment 1: Effects of dispersal status and host plant quality on individual-level performance

Mites randomly sampled from available test patches (i.e. mated 3 to 4 days old adult females) were placed individually on freshly cut 4 cm^2^ leaf squares placed on wet cotton and surrounded by moist tissue paper. Leaves were either stressful (4-week-old tomato, Solanum lycopersicum cv. Moneymaker) or suitable (bean). Forty replicates were made per dispersal status × host combination (*N_total_* = 160). We used population size (mites of all stages + eggs) after 10 days as a proxy of mite performance (Wybouw et al., 2015). Similar results were obtained using the number of eggs laid in the first 24 h as an alternative performance index (Supplementary Figure 1; correlation with 10-day population size, r = 0.65). We evaluated feeding rate by taking pictures of leaves after 24h (before any offspring of focal mites began to feed). Pictures were compared to those made at the start of the experiments, and the surface covered by feeding damage (chlorotic marks) was measured using ImageJ (Abramoff et al., 2004). We used the same images to measure average egg size and mite body length; two mites were not measured due to bad positioning at the time of the photograph.

### Experiment 2: Effects of dispersal status and host plant quality on changes in metabolic profiles

We placed dispersers or residents randomly sampled from patches - 3 to 4 days old mated females - by groups of 50 on fresh 4 cm^2^ leaf squares prepared as above. Between 4 and 6 replicates were made per dispersal × host treatment combination (*N_total_* = 20 samples). We left mites on leaves for 24 h, as preliminary tests showed that longer exposures to tomato at this density led to high (>20%) mortality. Mites were then collected in microtubes, weighted by group to the nearest 0.01 mg (mass_microtube+mites_ - mass_microtube_), and snap-frozen in liquid nitrogen before storage at −20 °C.

We then prepared samples for metabolomics analyses by gas chromatography–mass spectrometry (GC-MS), as described in Khodayari et al. (2013) and van Petegem et al. (2016). Samples were first homogenized into 300 µL of ice-cold 2:1 methanol-chloroform using a tungsten bead beating apparatus (RetschTM MM301; Retsch GmbH, Haan, Germany) at 25 Hz for 1.5 min. After addition of 200 µL of ice-cold ultrapure water, samples were centrifuged at 4000 g for 5 min at 4 °C. An aliquot of 240 μL of the upper aqueous phase (containing polar metabolites) was then transferred to new chromatographic glass vials. Samples were vacuum-dried (Speed Vac Concentrator, MiVac, Genevac Ltd., Ipswitch, England), and resuspended in 30 µL of 20 mg.L^−1^ methoxyamine hydrochloride (Sigma-Aldrich, St. Louis, MO) in pyridine, and kept under automatic orbital shaking at 40 °C for 60 min of incubation. Finally, 30 μL of N,O-Bis(trimethylsilyl)trifluoroacetamide (BSTFA; Sigma, CAS number 25561-30-2) was added, and the derivatization was conducted at 40 °C for 60 min under agitation. Prepared samples were then analyzed in a gas chromatography–mass spectrometry (GC-MS) system (Thermo Fischer Scientific Inc., Waltham, MA), using the same settings as in Khodayari et al. (2013) and van Petegem et al. (2016). The selective ion monitoring (SIM) mode was used to search for 60 primary metabolites common in arthropods and included in our spectral database (Van Petegem et al., 2016). Calibration curves were set up for each metabolite using pure reference compounds (1, 2, 5, 10, 20, 50, 100, 200, 500, 750, 1000, and 1500 μmol.L^−1^). Chromatograms were deconvoluted using XCalibur v2.0.7 (Thermo Fischer Scientific Inc.). Thirty-seven (out of 60) metabolites were successfully quantified in our samples and thus used in subsequent analyses. Concentrations were expressed in nmol per mg of mite fresh mass. Metabolites were grouped into six chemical categories based on the KEGG database hierarchy (Kanehisa et al., 2017): amines (N = 3), free amino acids (N = 14), organic acids (N = 5), polyols (N = 7), sugars (carbohydrates excluding polyols, N = 5), and other molecules (N = 3, gluconolactone, glycerol-3-phosphate, phosphoric acid).

### Statistical analyses

We analyzed the effects of dispersal status, post-trial host and their interaction using trait-specific models, fitted with R, version 3.5 (R Core Team, 2018).

For the first experiment, we used a quasi-Poisson generalized linear model for the number of mites at 10 days, and linear models for body size and egg volume (assuming spherical eggs). As 53.75 % of mites did not feed, leaf consumption was analyzed using a binomial-lognormal hurdle model (see e.g. Fletcher et al., 2005), with feeding probability analyzed as a binary variable (fed/did not feed, binomial GLM) and leaf consumption in mites that did feed (N = 60 on bean, 14 on tomato) using a Gaussian GLM with log link.

For the second experiment, we used a linear model to analyse mean fresh mass, and a permutational MANOVA (10000 permutations) as implemented in the `RRPP` package (Collyer and Adams, 2018) to analyze the multivariate set of metabolite concentrations. A classical parametric MANOVA could not be used due to the high-dimensional structure of our dataset (number of molecules > number of samples) (Collyer and Adams, 2018). Concentrations were scaled to unit standard deviation before analysis to avoid metabolites with high baseline concentrations having a disproportionate influence on the result (van den Berg et al., 2006).

Then, to determine whether dispersal status or host plant altered the concentrations of specific categories of metabolites, we used the generalized linear mixed model approach proposed by Jamil et al. (2013) to analyze plant community data. Indeed, as a one-to-one correspondence can be drawn between the structure of community ecology and metabolomics datasets (sites = replicates; species = molecules, species traits = molecular characteristics), methods designed in one context can be readily applied to the other. We analyzed the concentration of all molecules grouped in a common univariate vector (37 molecules × 20 replicates = 740 concentration records), as a function of metabolite categories, sample dispersal status, host plant and all interactions. Furthermore, we included random effects of replicate (intercept-only), to account for the non-independence of molecular responses from a same sample, as well as random effects of individual metabolite to account for molecule-specific responses (random intercept as well as dispersal, host, and dispersal x host effects). We fitted the model with the `glmmADMB` package (Fournier et al., 2012; Skaug et al., 2016), using the Gamma family (as concentrations are always continuous and > 0) and a log link. As the focus is on relative changes in concentrations, we used relative concentrations (mean concentration for all molecules set to 1) to avoid violations of model assumptions due to a few molecules present in much higher abundance than the others. We followed Schielzeth (2010)’s guidelines to make model coefficients directly interpretable even in the presence of higher-order interactions: dispersal status and host were included in the model as centered dummy variables (Resident or Bean = −0.5, Disperser or Tomato = 0.5). This and the log link lead to treatment × metabolite category model coefficients being directly interpretable as log(fold changes) in response to treatment. We estimated conditional and marginal R^2^ following Nakagawa et al. (2017), using the trigamma method. Plots were generated using the `ggplot2` and `cowplot` packages (Wickham, 2016; Wilke, 2019).

## Results

### Experiment 1: Effects of dispersal status and host plant quality on individual-level performance

Body size did not vary according to dispersal status and host plant, or their interaction (*ANOVA*, Table 1; mean length ± SE = 437 ± 2 μm). As expected from a putative stress treatment, mites exposed to tomato after dispersal trials performed worse than mites kept on bean (2.19 ± 0.57 vs 48.71 ± 7.69 individuals per leaf at 10 days, Table 1, Fig. 1), and were less likely to feed (Fig. 2; Table 1), with no significant effect of dispersal status or dispersal × host interaction in both cases (Table 1). Although feeding probability itself was not linked to dispersal status, dispersers that did attack their host plant fed less (26.99 ± 10.62 % less chlorotic damage) than residents (Fig. 2, Table 1).

**Figure 1.**
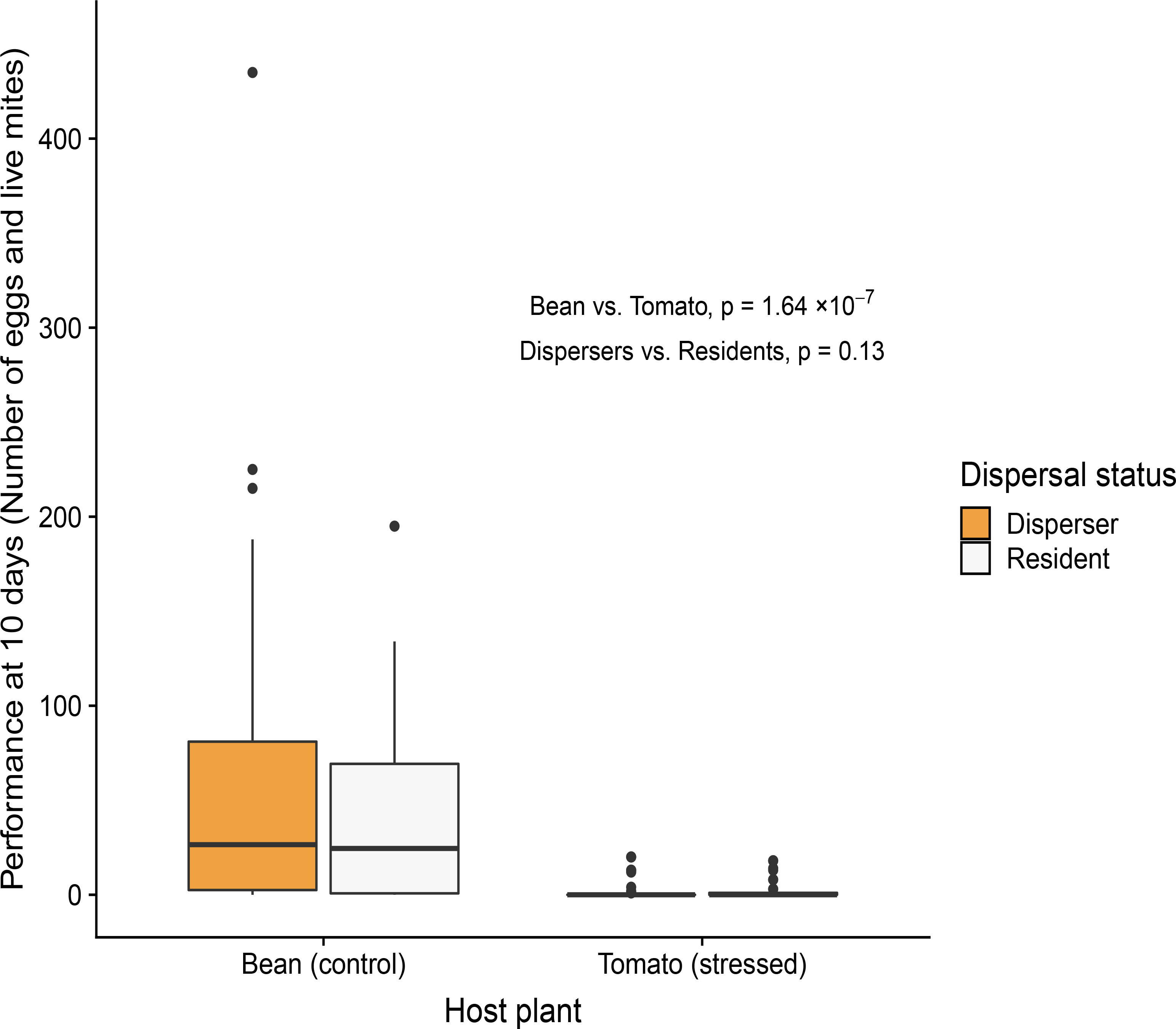
Effect of dispersal status and host plant on mite performance, measured as the number of live individuals (all stages, including eggs) present on leaves after 10 days in Experiment 1. See Table 1 for test details.

**Figure 2.**
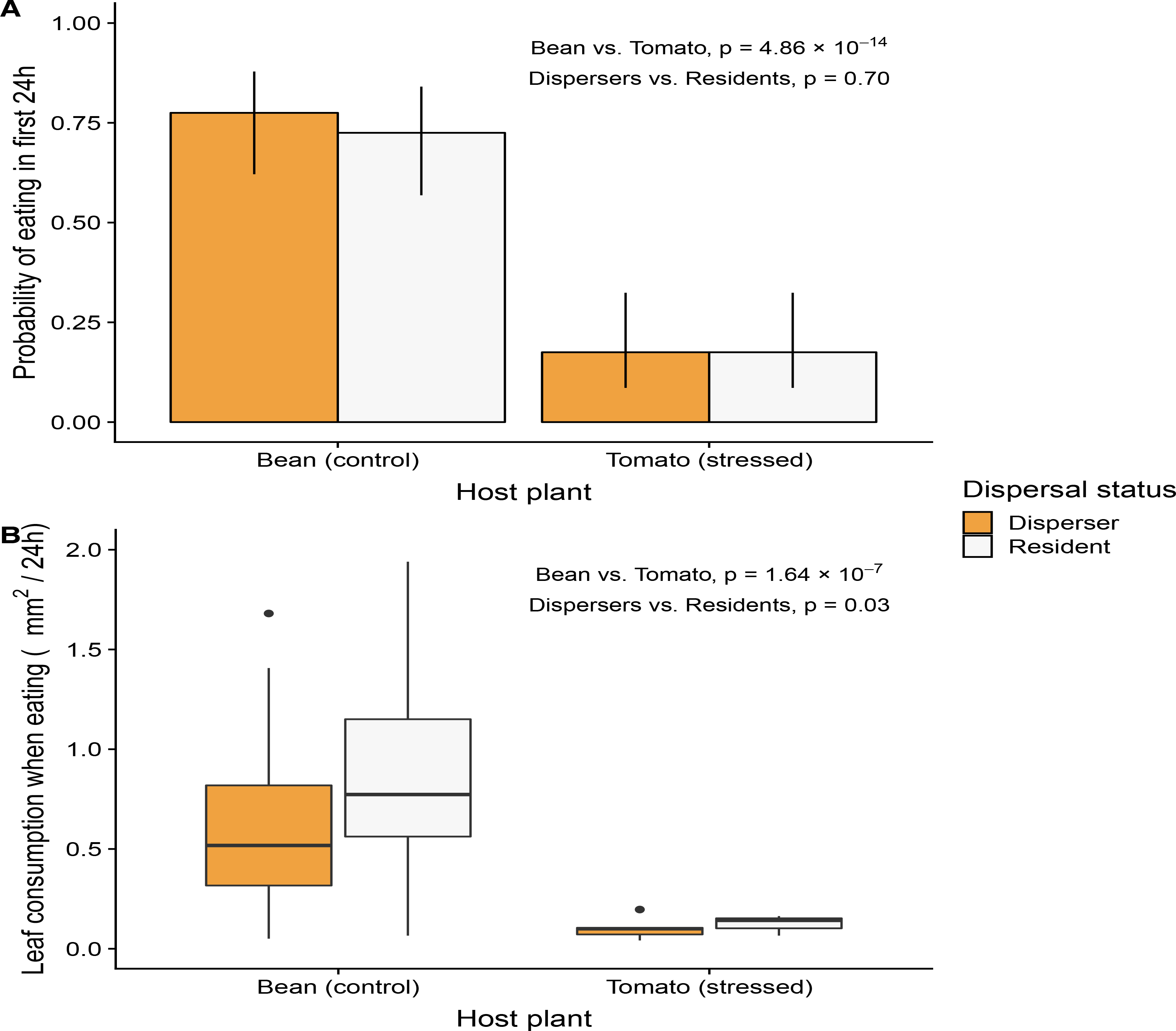
Effect of dispersal status and host plant on (A) spider mite feeding probability per 24h with 95% CI and (B) feeding intensity (leaf damage per 24h) in the 74 mites that did feed during the first 24 h of Experiment 1. See Table 1 for test details.

**Table 1.** Test statistics for the effect of dispersal status, host plant and their interaction on several *Tetranychus urticae* traits

| Trait |  | Model | Number of replicates | Dispersal status | Post-trial host plant | Dispersal x Host interaction |
| --- | --- | --- | --- | --- | --- | --- |
| <u>Experiment 1:</u> |  |  |  |  |  |  |
| Individual body size (length, $\mu\text{m}$) | | linear model | 158 | $F_{1,154} = 3.46; p = 0.06$ | $F_{1,154} = 2.66; p = 0.10$ | $F_{1,154} = 0.00; p = 0.96$ |
| Total population size at 10 days | | GLM (quasi Poisson) | 160 | $\chi^2 = 2.32, \text{df} = 1, p = 0.13$ | $\chi^2 = 81.97, \text{df} = 1, p < 2 \times 10^{-16}$ | $\chi^2 = 0.13, \text{df} = 1, p = 0.72$ |
| Feeding during the first 24h | Feeding probability | GLM (binomial) | 160 | $\chi^2 = 0.15, \text{df} = 1, p = 0.70$ | $\chi^2 = 56.79, \text{df} = 1, p = 4.86 \times 10^{-14}$ | $\chi^2 = 0.12, \text{df} = 1, p = 0.73$ |
| | Feeding intensity (chlorotic damage, $\text{mm}^2$) | GLM (log-normal) | 74 | $\chi^2 = 4.83, \text{df} = 1, p = 0.03$ | $\chi^2 = 27.42, \text{df} = 1, p = 1.64 \times 10^{-7}$ | $\chi^2 = 0.00, \text{df} = 1, p = 0.97$ |
| Volume of average egg laid in the first 24h ($\mu\text{m}^3$) | | linear model | 136 | $F_{1,132} = 6.00; p = 0.02$ | $F_{1,132} = 17.51; p = 5.18 \times 10^{-5}$ | $F_{1,132} = 0.16; p = 0.69$ |
| <u>Experiment 2:</u> |  |  |  |  |  |  |
| Individual body mass (average of 50 individuals per replicate, $\mu\text{g}$) | | linear model | 20 | $F_{1,16} = 1.98; p = 0.63$ | $F_{1,16} = 11.32; p = 0.25$ | $F_{1,16} = 0.01; p = 0.93$ |
| Multivariate metabolic profile (37 molecules) | | RRPP- Permutational MANOVA | 20 | $F_{1,16} = 2.10, p = 0.03$ | $F_{1,16} = 2.33, p = 0.02$ | $F_{1,16} = 1.68, p = 0.06$ |

Dispersers laid smaller eggs than residents (9.39 ± 0.19 vs 10.04 ± 0.19 ×10^5^ μm^3^), and mites exposed to tomato larger eggs than mites kept on bean (10.27 ± 0.18 vs 9.16 ± 0.19 ×10^5^ μm^3^; Fig. 3, Table 1). There was again no significant dispersal × host interaction.

**Figure 3.**
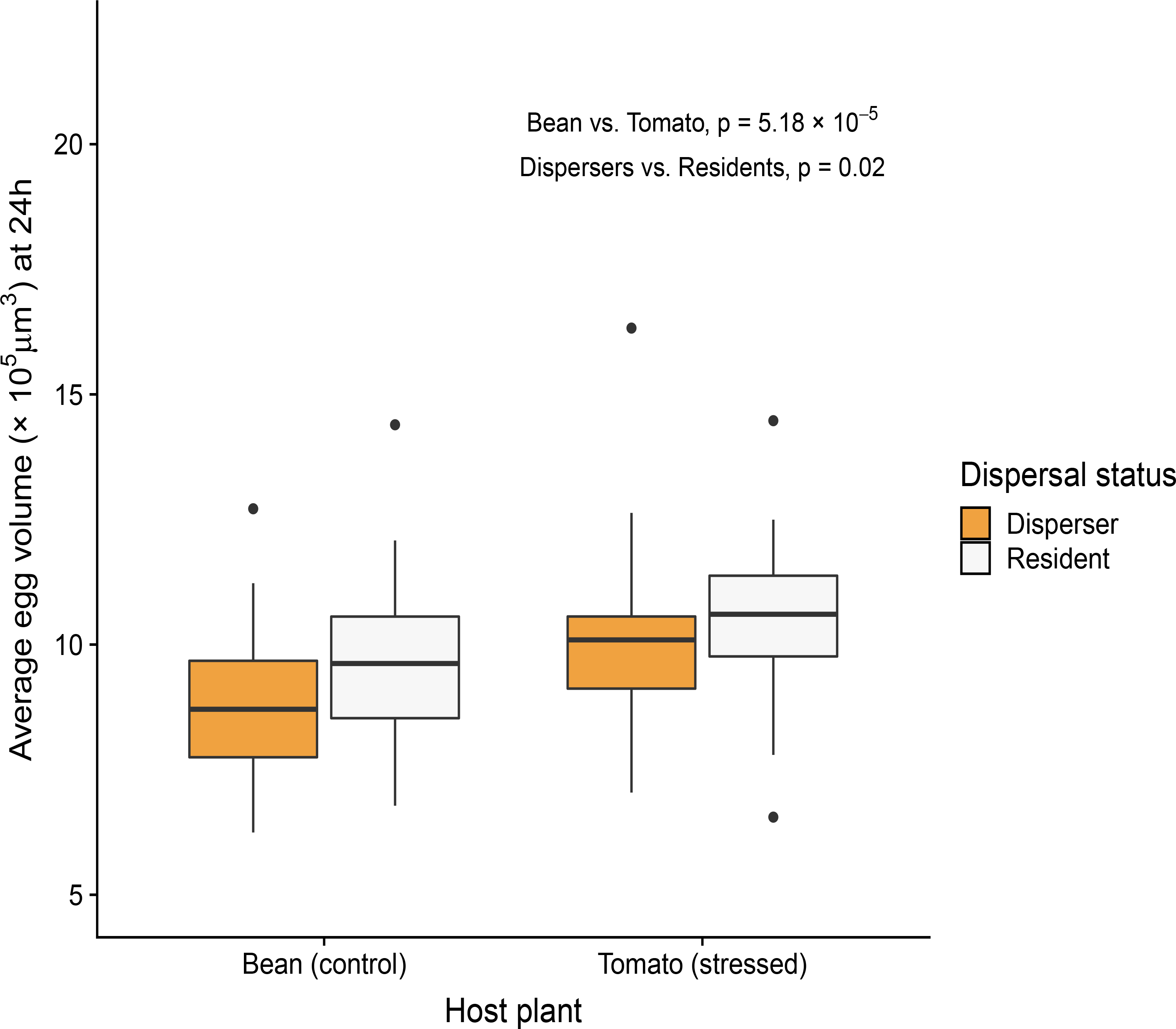
Mean egg volume per mother as a function of dispersal status and host plant (for the 138 mites that laid eggs in the first 24 h of Experiment 1). See Table 1 for test details.

### Experiment 2: Effects of dispersal status and host plant quality on changes in metabolic profiles

Mean mite fresh mass was independent from dispersal status and host plant, or their interaction (*ANOVA*, Table 1; mean individual mass ± SE = 12 ± 1 μg). The multivariate metabolic profile was significantly influenced by both dispersal status and host treatment, but with no significant interaction among these terms (Table 1). The molecular category-level GLMM explained 52.36 % of the total variation in concentrations (*R*^2^_*m*_) with 13.37 % explained by fixed effects (inter-category variations and category × treatment effects, *R*^2^_*c*_). The model successfully predicted mean relative concentrations for each treatment × molecular category (Pearson correlation between predicted and observed values = 0.95) and for each treatment × molecule (*r* = 0.97) combinations (see Supplementary Figure 2). Metabolite mean relative concentrations were significantly influenced by dispersal status and host plant, but not by their interaction (see Table 2). Analysis of model coefficients indicates that residents have on average higher concentrations of amino acids and amines than dispersers (Table 2, Fig. 4). Tomato-exposed mites on the other hand have on average lower concentrations of sugars than bean-kept controls (Table 2, Fig. 4).

**Table 2.** Model coefficients for the generalized linear mixed model (Gamma family, log link) explaining metabolite relative concentration as a function of dispersal, host plant and metabolite molecular category. Fixed effects are given with their SE; bold values are significantly different from zero based on 95% confidence intervals. The model was re-parametrized following Schielzeth (2010) to facilitate interpretation. First, the global intercept was removed and replaced by category-specific intercepts. Second, Dispersal and Host variables were centered so their coefficients could be interpretable directly even in the presence of interaction (Resident or Bean = −0.5, Disperser or Tomato = 0.5).

| Fixed effects coefficients ± SE | | | | |
| --- | --- | --- | --- | --- |
| Molecular category | Category Intercept | Category × Dispersal status | Category × Host | Category × Dispersal × Host |
| Amines | -0.20 ± 0.16 | **-0.60 ± 0.30** | 0.45 ± 0.29 | -0.17 ± 0.69 |
| Amino acids | -0.17 ± 0.10 | **-0.41 ± 0.18** | -0.30 ± 0.18 | 0.23 ± 0.40 |
| Carbohydrates (excluding polyols) | **-0.32 ± 0.13** | 0.03 ± 0.25 | **-0.49 ± 0.24** | -0.44 ± 0.56 |
| Organic acids | -0.07 ± 0.13 | -0.05 ± 0.25 | 0.05 ± 0.24 | -0.41 ± 0.56 |
| Polyols | -0.01 ± 0.12 | 0.18 ± 0.22 | -0.02 ± 0.21 | -0.43 ± 0.49 |
| Others | -0.20 ± 0.16 | -0.37 ± 0.30 | -0.55 ± 0.29 | -0.56 ± 0.69 |
| Random effects variances |  |  |  |  |
| Molecule Intercept | 0.05 |  |  |  |
| Molecule × Dispersal status | 0.16 | | | |
| Molecule × Host plant | 0.14 | | | |
| Molecule × Dispersal status × Host plant | 0.99 | | | |
| Sample Intercept | 0.08 |  |  |  |

**Figure 4.**
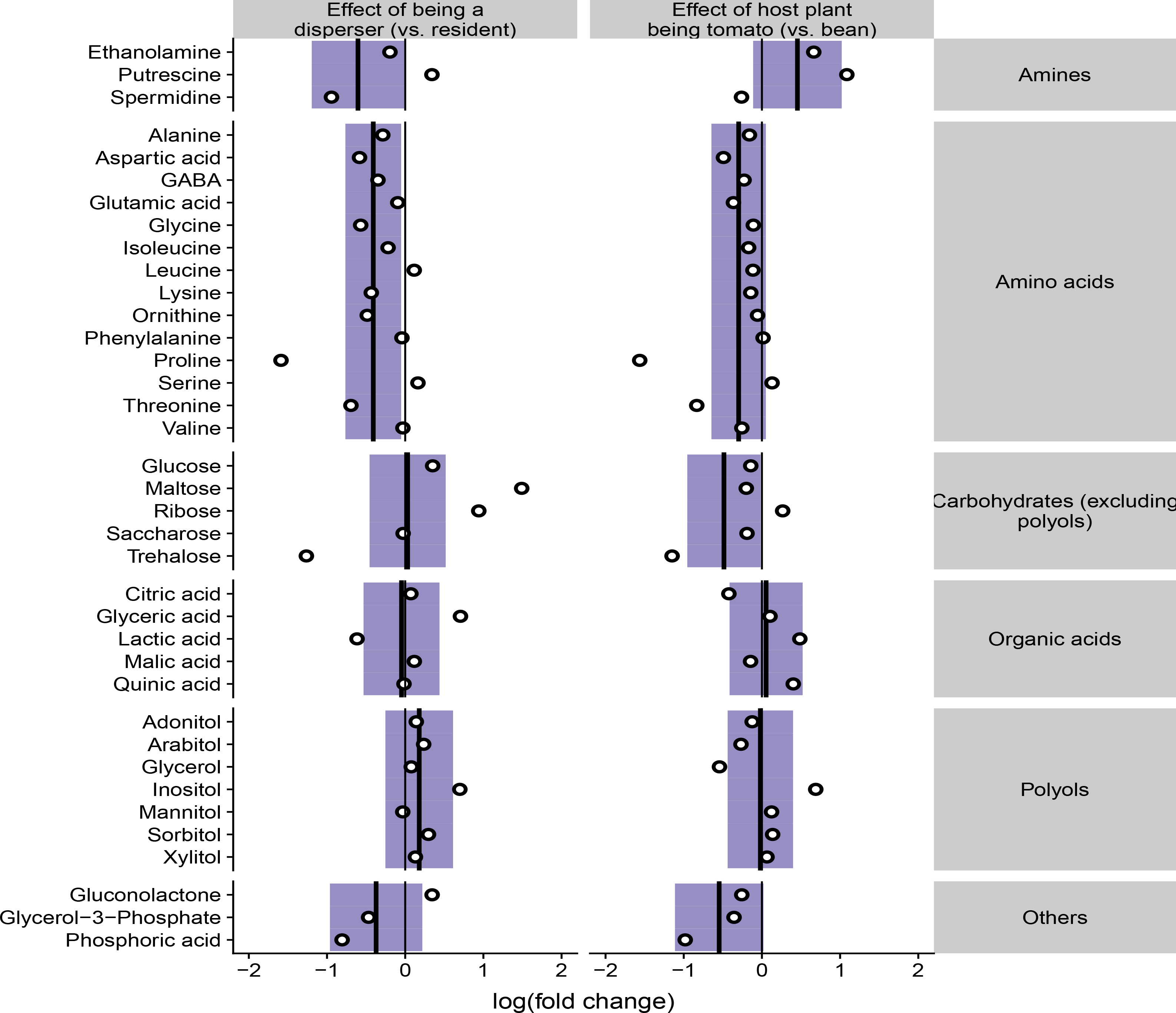
Effect of dispersal status and host plant on relative metabolite concentration in Experiment 2 (Table 2). Molecules are grouped by “categories” based on KEGG (Kanehisa et al., 2017); colored rectangles denote 95% confidence bands for the category-level effect of dispersal/host, bold segments the mean predicted effect. Observed mean log(fold changes) for each molecule are presented as white-filled dots for illustrative purposes.

## Discussion

By combining dispersal assays, “classical” phenotypic trait measurements and metabolomics, we developed an unprecedented multi-level study on dispersal syndromes and the way they may be shaped by environmental context (Bonte and Dahirel, 2017). We found that dispersers and residents of the spider mite *Tetranychus urticae* differed in several traits forming a newly described dispersal phenotypic syndrome for this species, including metabolic profiles rarely investigated in that context. However, contrary to our hypotheses and despite clear costs of biotic stress exposure, phenotypic differences between dispersers and residents were not influenced by host plant. This independence of dispersal syndrome and stress response is maintained across a wide range of traits, including life-history, physiological and feeding traits. This is especially surprising as exposure to a stressful host had by itself major consequences for mite phenotype, influencing all tested traits (Table 1, except adult body size, which was reached prior to exposure), including traits involved in the dispersal syndrome. This suggests that maintaining fitness prospects by resisting stress or by escaping unfavourable habitats are here unrelated strategies that may be selected independently.

Mites from a bean-adapted lineage performed worse and fed much less when left on tomato compared to bean controls (Figs 1, 2). Most tomato-exposed mites even stopped feeding altogether (Fig. 2). Additionally, metabolic profiles of tomato-exposed mites were specifically characterized by lower concentrations of sugars (Fig. 4). This is a physiological signature of a starved state, as individuals then consume their carbohydrate reserves (e.g. Dus et al., 2011; Shi et al., 2017). Our results are here in line with previous studies. Indeed, although *Tetranychus urticae* is overall a generalist species and resistance to a given stressful host evolves readily (within a few generations), tomato is a consistently challenging host for unacclimated or unadapted mites (Alzate et al., 2017; Marinosci et al., 2015; Wybouw et al., 2015), possibly because of its content in toxic secondary metabolites (Wybouw et al., 2015). However, we found evidence that *Tetranychus urticae* is able to partly plastically adjust to such stressful conditions. Mites laid fewer eggs on tomato compared to bean, but they laid larger eggs (Fig. 3). In *Tetranychus urticae*, egg size has marked and direct consequences for fitness: larger eggs are more likely to survive to maturity, and yield on average larger adults, sex being equal (Macke et al., 2011). In this haplodiploid species, larger eggs are also more likely to be fertilized, and hence female (Macke et al., 2012). Our results here line up with a previous study showing that female *Tetranychus urticae* having experienced, or experiencing poor dietary environments are more likely to lay female eggs (Wrensch and Young, 1983). Female larvae are more likely than males to survive food stress during development (Wrensch and Young, 1983). Laying fewer but larger eggs on tomato may thus be a plastic response increasing mite initial success chances on a challenging host plant, and thus the odds of eventual successful genetic adaptation (Crispo, 2007; West-Eberhard, 2003). Note that, with our experimental design, we unfortunately cannot disentangle here these three consequences of host plant induced egg size changes (higher survival, larger size, altered sex ratio) on the second generation. Indeed, we did not raise eggs in isolation, and treatments with larger eggs also had lower population density after egg hatching (contrary to e.g. Macke et al., 2011). Nonetheless, the various consequences of tomato exposure for mite physiology, feeding and life-history are all compatible with plastic responses to food stress induced by feeding deterrence, possibly associated with direct toxicity of some tomato secondary metabolites.

We found that dispersers and residents differed in a suite of physiological, life-history and performance traits, providing evidence of a complex multivariate dispersal syndrome in *Tetranychus urticae*. Dispersers and residents had different metabolic profiles, mostly due to the former having on average lower concentrations in amino acids than the latter (Fig. 4, Supplementary Figure 2). Given similar resource quality, such differences may arise through two non-exclusive mechanisms: differences in amino acid uptake from the host, and/or differences in amino acid production or degradation by mites themselves. While proof of the latter can only come from deeper physiological studies, our results support at least a partial role of the former mechanism: although dispersers were as likely to feed as residents, dispersers that did feed extracted fewer resources from their host plant per unit of time, based on the lower amount of damage dealt (Fig. 2). Nitrogen/amino acids availability plays a major role in T. urticae fecundity and performance (Nachappa et al., 2013; Wermelinger et al., 1985). Accordingly, although dispersers and residents had the same number of offspring (Fig. 1, Supplementary Figure 1), dispersers’ eggs were notably smaller than residents’ (Fig. 3). As mentioned above, egg size has major impacts on offspring future survival and success (Macke et al., 2011; Potter et al., 1976), so adults developed from smaller dispersers’ eggs will likely be less competitive against offspring of residents. Smaller eggs in dispersers may also result from sex-ratio adjustments by egg-laying females towards more males (Macke et al., 2011; Macke et al., 2012), but we believe it is unlikely in the present case. Indeed, if we assume dispersing females are more likely than residents to find themselves in a newly founded population with few to no other females, then they should produce fewer males, and not more, based on Local Mate Competition theory (Macke et al., 2012).

Altogether, the observed syndrome linking dispersal to feeding, life history and metabolic profile is consistent with the hypothesis of a trade-off between dispersal and foraging efficiency proposed by Fronhofer and Altermatt (2015), to explain population and trait dynamics observed during experimental range expansions. In addition to explaining our present results, the existence of this trade-off in *T. urticae* provides a mechanism for the documented fitness costs incurred by dispersers forced to stay in densely populated/ resource limited contexts (Bonte et al., 2014), i.e. in contexts where they would be outcompeted by more “voracious” residents. By contrast, “prudent” use of resources by dispersers would be advantageous in newly colonized habitats where and competition by faster eaters is limited (Bonte et al., 2014; Fronhofer and Altermatt, 2015).

Phenotypic divergence between dispersers and residents was however not total, as they were impossible to differentiate on several tested axes of phenotypic trait variation. There was no difference in body size (length or mass) between dispersers and residents, despite evidence that larger individuals have competitive advantages in mites, including *Tetranychus urticae* (Potter et al., 1976; Walzer and Schausberger, 2013), and that body size influences dispersal tendency and/or success in other species (e.g. Moore et al., 2006; O’Sullivan et al., 2014). While we recorded differences in egg size (see above, Fig. 3), dispersers did not differ from residents in terms of performance at 10 days, hinting at limited differences in egg and larval survival (Fig. 1, see also Supplementary Figure 1 for fecundity at 24h). Our findings in our one generation experiment are here similar to those of Tung et al. (2018) in *Drosophila melanogaster* after several generations of selection for dispersal. They suggested that this stability of body size and fecundity despite the constitutive and induced costs of dispersal was offset by a shorter lifespan; our experiment was not designed to test this hypothesis. An alternative and non-exclusive explanation for the absence of size- or performance-syndromes is based on the fact that we only assayed mite life-history traits under low-density conditions in the present study. Indeed, Bonte et al. (2014) showed that dispersers only fared worse than residents in competitive, high-density contexts. We therefore consider it possible that syndromes linking dispersal to body size and fecundity may only be detectable when accounting for plasticity across a range of environmental variation (Bonte and Dahirel, 2017).

Dispersal syndromes were however independent from stress exposure, a result that was consistent across life history traits and metabolic profiling: overall, a mite dispersal status had no influence on the intensity of its later phenotypic response and tolerance to tomato. This was contrary to our expectations based on hypothesized dispersal/stress tolerance trade-offs and the results of previous meta-population level evolutionary studies (De Roissart et al., 2015; De Roissart et al., 2016), that showed that spatial contexts selecting for decreased dispersal also selected for higher stress tolerance. Our results show that that correlated response does not result from pre-existing correlations between these two strategies to mitigate the effects of unfavorable conditions. Dispersal and stress responses are actually independent down to their physiological aspects, which involve shifts in different categories of metabolites (sugars in response to stress versus amino acids in dispersers, Fig. 4). In that context, the apparent population-level correlation between the two traits in evolutionary experiments may result from independent responses to the same selective constraint (Ronce and Clobert, 2012), rather than from pre-existing physiological trade-offs or genetic constraints. For instance, demographic and environmental conditions experienced before dispersal and during development may simultaneously shape dispersal and stress tolerance (Baines and McCauley, 2018; Endriss et al., 2018; Henry et al., 2018; Kent et al., 2009). This further confirms that syndromes are not necessarily fixed and may evolve rapidly within species, and that this flexibility needs to be accounted for to fully understand spatial and population dynamics (Bonte and Dahirel, 2017). Complicating the picture is the fact that, in spatially heterogeneous contexts where dispersal is relevant, individuals may experience variation in not one, but several biotic and abiotic stressors simultaneously or in close succession (Fronhofer et al., 2018). Because of this, and as more research show that the ecological impacts of stressors can be non-additive (Côté et al., 2016), future studies on the physiological underpinnings of dispersal-stress syndromes should aim to study ecologically relevant combinations of stressors. Such research would fundamentally increase our ability to understand the consequences of stress for movement and forecast its implications for the meta-population dynamics of threatened, range-shifting or pest species in a changing world.

## Author contributions

Conceptualization: MD, SM, DB; Methodology: MD, SM, DR, DB; Software: MD, SM; Validation: MD; Formal analysis: MD; Investigation: MD, SM, DR; Resources: MD, SM, DR, DB; Data curation: MD; Writing – original draft preparation: MD; Writing – review and editing: MD, SM, DR, DB; Visualization: MD; Supervision: DB; Project administration: DB; Funding acquisition: DB, DR

### Data accessibility

Data are available on Figshare (DOI: 10.6084/m9.figshare.7857260)

### Funding

This project was funded by the Research Foundation – Flanders (FWO) (G.018017N). DB was further supported by the FWO research network EVENET (grant W0.003.16N), DR by the CNRS (ENVIROMICS call) and the Institut Universitaire de France. SM is a holder of a FWO doctoral fellowship, and MD held a Fyssen Foundation postdoctoral fellowship at the time of the research.

### Ethical statement

The study complies with all relevant national and international ethical laws and guidelines. No ethical board recommendation was needed to work on *Tetranychus urticae*.

