## Supplementary Figures for "The distinct phenotypic signatures of dispersal and stress in an arthropod model: from physiology to life history"

**Supplementary Figure 1.** Effect of dispersal status and host plant on fecundity, measured as the number of eggs laid in the first 24h of Experiment 1. Dispersal effect: $X^2 = 1.47$, df = 1, p = 0.23 ; Host plant effect: $X^2 = 43.34$, df = 1, $p = 4.59 \times 10^{-11}$; Interaction effect: $X^2 = 0.00$, df = 1, p = 0.95 (quasi-Poisson GLM).

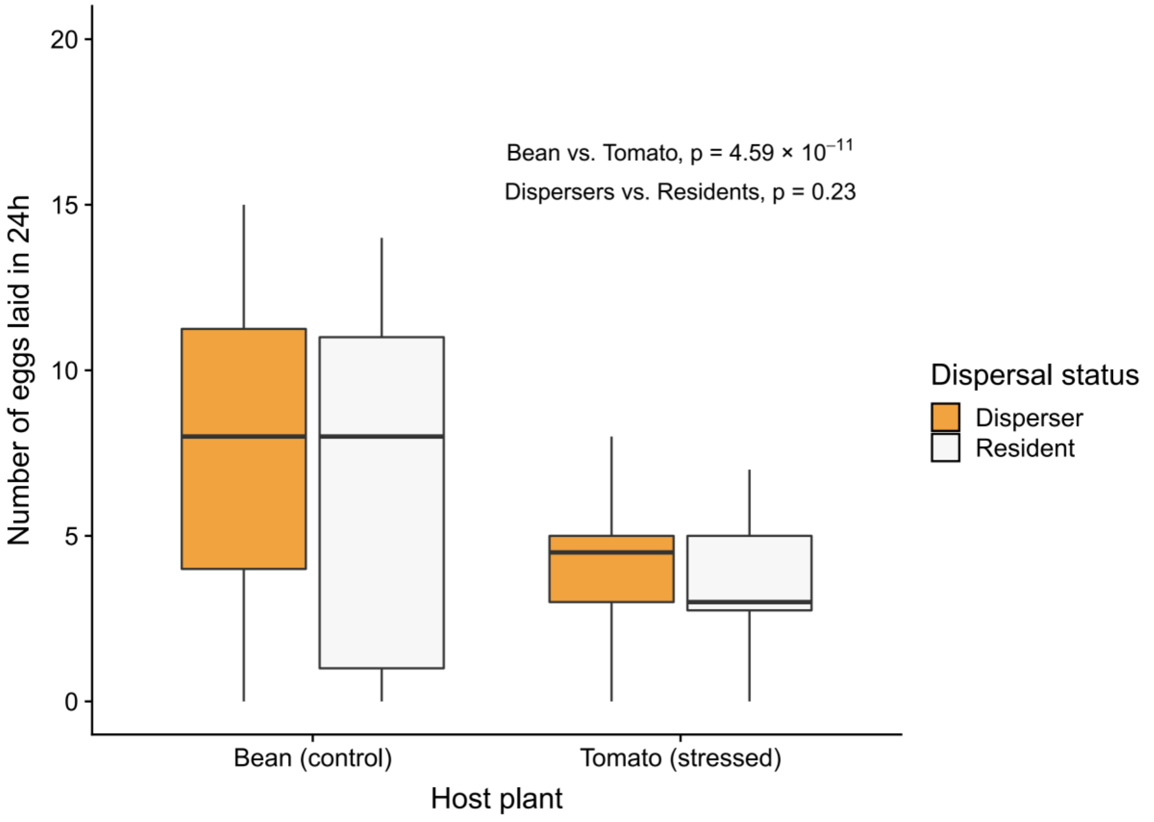


**Supplementary Figure 2. (next pages)** Observed metabolite concentrations in Experiment 2. Box and black dots: observed values, coloured dots: predicted means based on fixed and random effects of the model presented Table 2 of the main text. Predictions are based on relative concentrations (mean value = 1) and back-transformed to actual concentrations using observed means. Molecular categories are presented in the same order as in Figure 4 (main text). Number of replicates = 5, 6, 5 and 4 for Dispersers on Bean, Residents on Bean, Dispersers on Tomato and Residents on Tomato, respectively.

**Supplementary figures -** The distinct phenotypic signatures of dispersal and stress in an arthropod model: from physiology to life history

Maxime Dahirel, Stefano Masier, David Renault, Dries Bonte

**Supplementary Figure 2. (continued)**

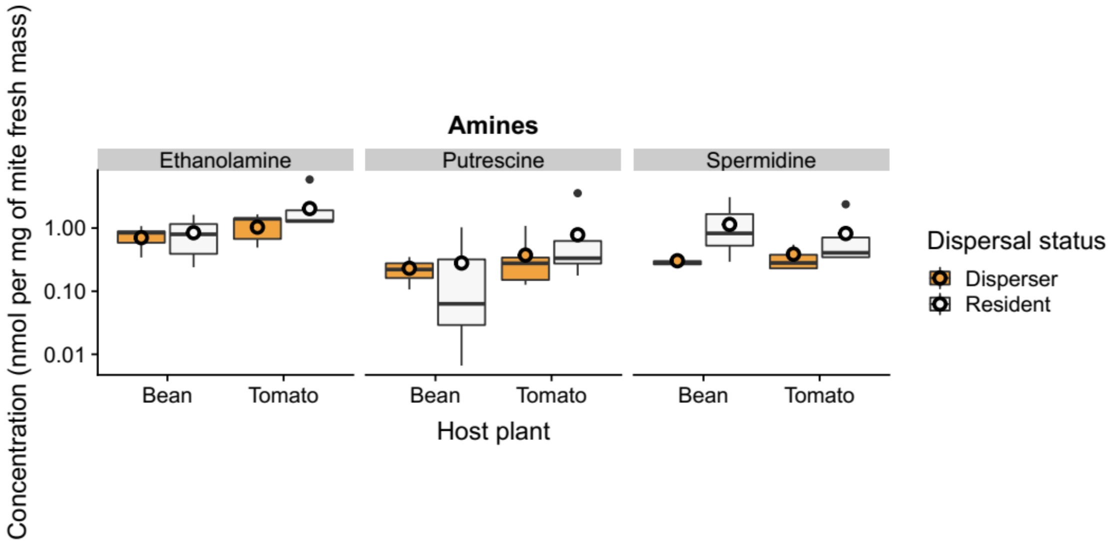

**Supplementary figures -** The distinct phenotypic signatures of dispersal and stress in an arthropod model: from physiology to life history

Maxime Dahirel, Stefano Masier, David Renault, Dries Bonte

**Supplementary Figure 2. (continued)**

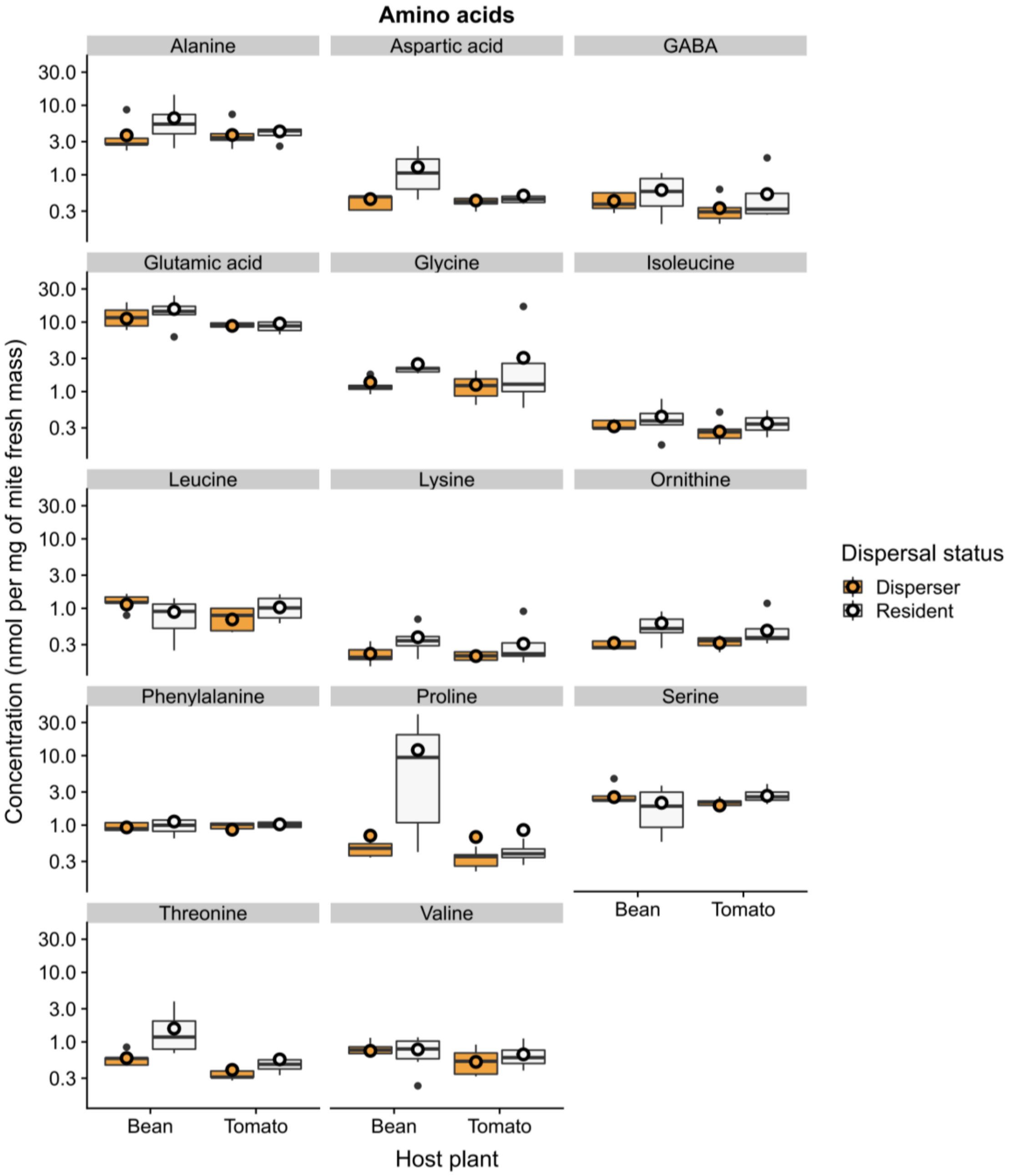

**Supplementary figures -** The distinct phenotypic signatures of dispersal and stress in an arthropod model: from physiology to life history

Maxime Dahirel, Stefano Masier, David Renault, Dries Bonte

**Supplementary Figure 2. (continued)**

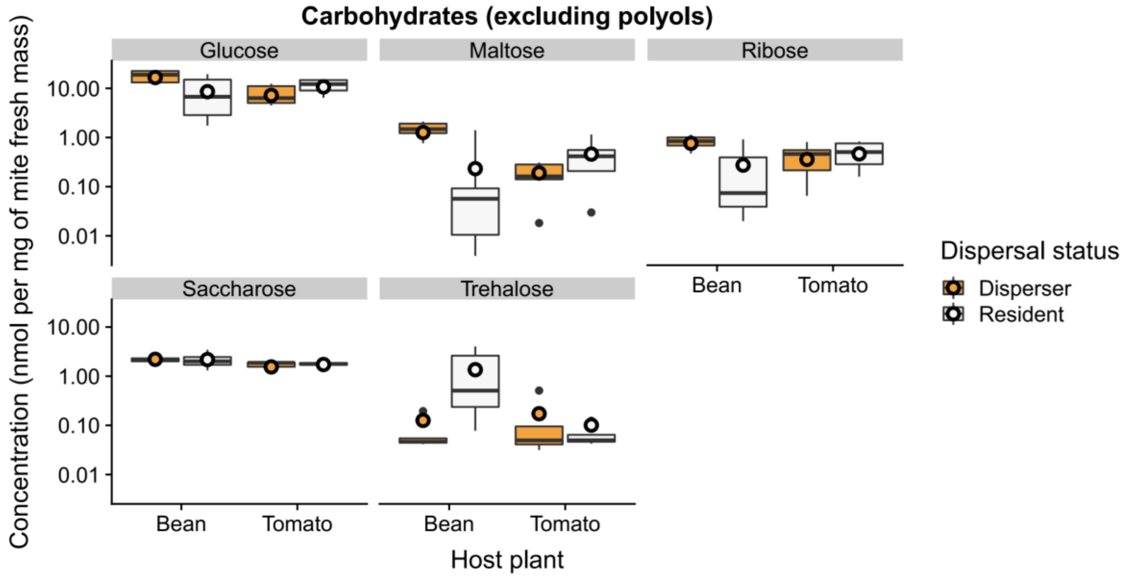


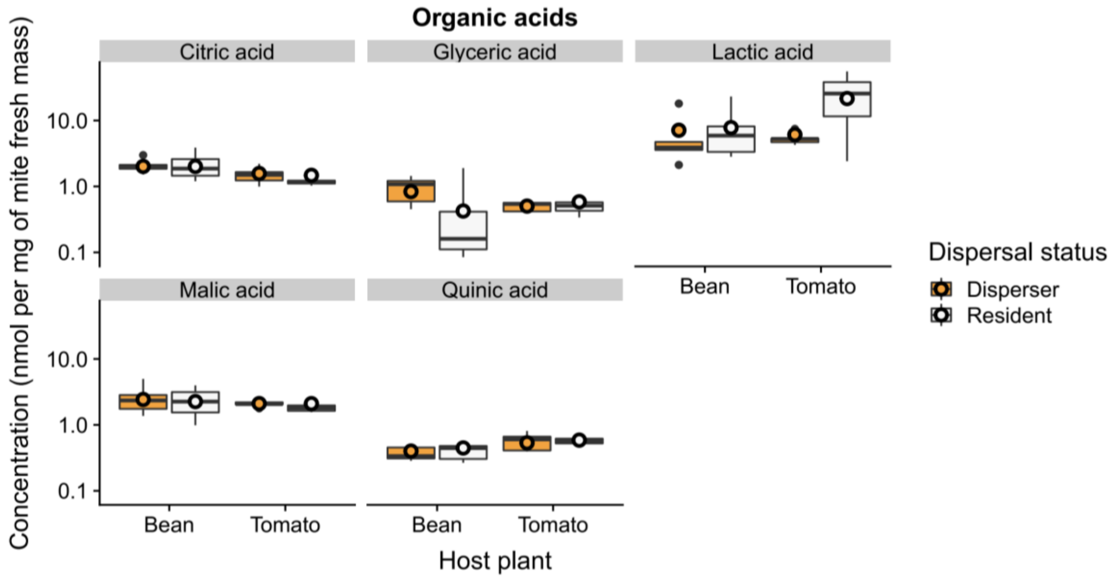

**Supplementary figures -** The distinct phenotypic signatures of dispersal and stress in an arthropod model: from physiology to life history

Maxime Dahirel, Stefano Masier, David Renault, Dries Bonte

**Supplementary Figure 2. (end)**

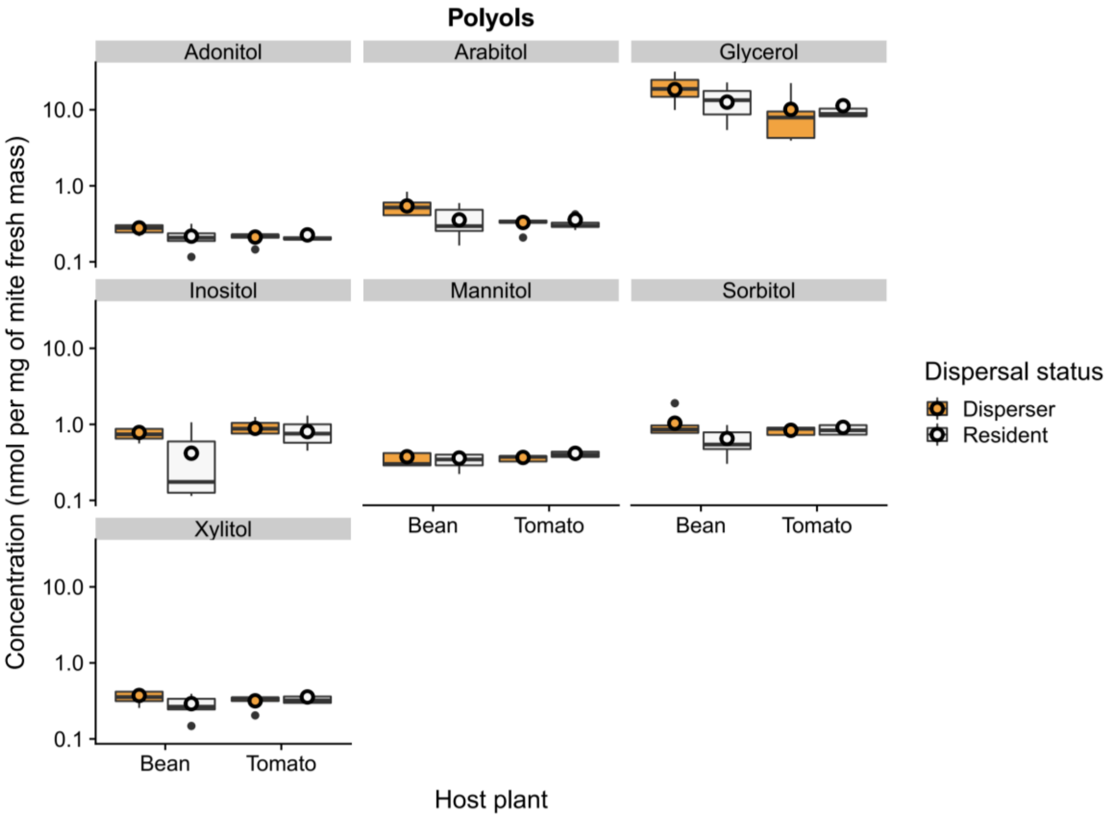


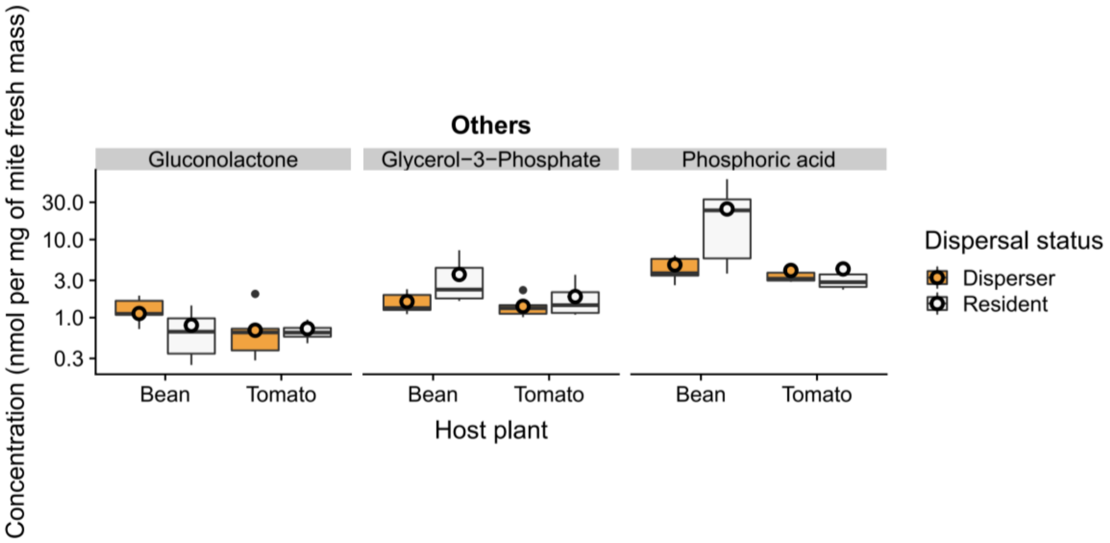
